# Impact of Image Representation on Deep Learning-Based Single-Cell Classification by Holographic Imaging Flow Cytometry

**DOI:** 10.64898/2026.02.26.708207

**Authors:** Daniele Pirone, Beatrice Cavina, Giusy Giugliano, Francesca Nanetti, Francesca Reggiani, Lisa Miccio, Ivana Kurelac, Pietro Ferraro, Pasquale Memmolo

## Abstract

Accurate cell type classification is essential for a wide range of biomedical applications, including disease diagnosis, drug discovery, and the study of cellular processes. Holographic imaging flow cytometry (HIFC) provides label-free quantitative phase imaging (QPI) of individual cells, enabling classification based on phase images. However, reconstructing holograms into phase images involves multi-step image processing, which introduces substantial computational overhead. The availability of diverse image representations across holographic reconstruction stages allows for flexible analytical strategies, enabling the optimization of trade-off between classification accuracy and computational efficiency. Moreover, deep learning offers an efficient alternative, accelerating the reconstruction process while performing accurate classification. However, despite its importance, this optimization challenge remains largely unexplored in the current literature. Here, we present the first systematic evaluation aimed at balancing classification accuracy with computational efficiency, highlighting how different image representations affect overall performance. We focus on a binary classification task discriminating natural killer cells from breast cancer cells. Six distinct classification pipelines are evaluated: direct processing of raw holograms, analysis of demodulated complex fields (CFs), refocused CFs, unwrapped phase images, and two deep learning-based methods that either replace the automatic refocusing stage or perform end-to-end hologram-to-phase reconstruction. For each strategy, we assess both computational cost and classification performance. Our results reveal a clear trade-off: reconstructed phase images provide the highest accuracy, whereas simpler representations or accelerated reconstruction methods significantly reduce processing time with minimal loss of accuracy. A Pareto analysis identifies the optimal set of strategies, offering practical guidelines for selecting image representations and processing pipelines based on available hardware and desired performance. Thus, this work offers a systematic framework for high-throughput deep learning classification in HIFC, serving as a potential reference for future biomedical applications.

## 1. Introduction

Accurate classification and/or identification of cell types is crucial in a wide range of life science and biomedical applications, including disease diagnosis, neoplasia subtype classification, understanding of cellular processes, and drug testing and discovery [1]. Traditional methods for single-cell inspection, such as conventional imaging (e.g., fluorescence microscopy), present significant limitations. Label-free microscopy techniques are preferable as they simplify sample preparation and preanalytical procedures by eliminating chemical treatments. Among these, quantitative phase imaging (QPI) [2,3] offers a valuable approach, encoding the distinctive fingerprint of each cell in variations of optical path length, determined by the combination of physical thickness and refractive index. However, expert identification of specific cells is hindered by the lack of specificity compared to fluorescence imaging [4]. In this context, Artificial Intelligence (AI) is transforming QPI for biomedical applications by enabling faster and more accurate phase retrieval and reconstruction, as well as advanced characterization and identification of cell populations [5–7]. AI-QPI increases imaging throughput, improves classification accuracy, and makes systems more accessible to nonexperts due to the advantages of label-free imaging [8]. Within supervised learning frameworks, where labels for each cell image are available, AI-QPI has been successfully applied to diverse classification problems. Notably, it has delivered significant results in the classification of different cancer cells [9–28], immune cells [29–33], red blood cells [34–40], cell life cycles [41,42], sperm cells [43–45], drug screening and discovery [46–48], and bacterial analysis [49–51].

Broadly, these contributions can be categorized into classical machine learning approaches, based on handcrafted feature extraction, and deep learning solutions, typically employing convolutional neural networks (CNNs). Machine learning-based methods achieve acceptable accuracies in many applications [9–18,29,30,34–36,41,43,46–48]. While standard morphological and texture features are commonly used, incorporating prior knowledge of distinctive biophysical cellular characteristics is often essential for high classification performance. Deep learning approaches, in contrast, have been introduced to meet higher accuracy requirements and tackle more challenging scenarios [19–28,31–33,37–40,42,44,45,49–51].

In the context of holographic imaging flow cytometry (HIFC), the possibility to acquire digital holograms of single-cells at high rate is essential to provide enough data for machine/deep learning tools. Some HIFC approaches even utilize raw holograms for classification rather than reconstructed phase images [20,22,26,28]. This is advantageous for high-speed analysis, as it eliminates the computational overhead of the holographic reconstruction pipeline, including zero-th and twin order filtering, refocusing, and phase unwrapping, thus dramatically reducing processing cost. However, raw holograms also encode distortions and noise [22], which are partially removed during reconstruction, thereby improving the prediction accuracy of learning algorithms. Consequently, finding the optimal balance between computation time (and available hardware) and classification accuracy remains a significant challenge.

In this paper, we address, for the first time to the best of our knowledge, the important challenge of identifying an optimal deep learning strategy in HIFC-based single-cell classification framework for balancing classification accuracy, processing and prediction time, as well as memory usage, thus quantifying the impact of image representations across holographic reconstruction steps. To validate our approach, we consider a binary classification problem, aiming to distinguish between MDA-MB-436 triple-negative breast cancer (TNBC) cell line and natural killer-derived NK92 cells. TNBC is a highly aggressive subtype of breast cancer showing poor prognosis and being resistant to conventional therapeutic approaches, including both chemotherapy and radiotherapy [52]. However, TNBC showed an unexpectedly higher response to immunotherapy-based protocols, such as the KEYNOTE522 [53,54], compared to other breast cancer subtypes. The proper stimulation of immune system drastically enhanced neoadjuvant chemotherapy response and patient overall survival. A recent study investigated the role of different immune cells in promoting chemotherapy efficacy in TNBC patients [55]. NKs were found enriched in the diagnostic biopsies of patients that showed complete response, suggesting it to be relevant in defining the therapy outcome. Thus, in the context of TNBC, we chose to analyze NKs which are emerging not only as potential prognostic biomarkers [56] but are being considered as a novel immunotherapy strategy [57]. In our analysis, we first evaluate classification performance across four different image representations fed to a neural network: (i) raw holograms, (ii) demodulated complex fields (CFs) obtained by removing zero-order and twin images, (iii) refocused CFs generated through automated refocusing reconstruction, and (iv) unwrapped phase images. In addition, to achieve an optimal balance between classification accuracy and computational efficiency, we explore deep learning-based approaches to accelerate the most time-consuming reconstruction stages. Specifically, we investigate CNN models capable of either (v) performing end-to-end phase reconstruction [58-64] or (vi) replacing only the automatic refocusing step [65-68], allowing us to significantly reduce processing time while maintaining high classification performance. We conduct a Pareto analysis of the six considered strategies, showing that an optimal choice can be identified. Our comprehensive analysis highlights that optimizing classification across holographic reconstruction stages requires balancing accuracy and computational cost, providing clear guidelines for selecting the most suitable image representation given available hardware and software resources. These results demonstrate that different strategies can be adopted depending on whether accuracy, speed, or computational efficiency is prioritized. Thus, we believe this study could offer a systematic framework for optimizing deep learning classification in HIFC and serve as a valuable reference for future research and biomedical applications.

## 2. Methods

### 2.1 Sample preparation

Natural killer-derived NK92 cell line (#CRL-2407) and MDA-MB-436 cell line (#HTB-130) were purchased from ATCC. All cell lines have been authenticated through SNP or STR profiling by Multiplexion Gmbh (Heidelberg, Germany). NK92 cell line was maintained in suspension and grown in Mem-alpha medium without nucleosides (Thermo Fisher #12561056) + 12.5% FBS South America (Gibco #A5256701) + 12.5% Horse serum (Thermo Fisher #26050088) + 1.5g/l Sodium Bicarbonate (Thermo Fisher #25080094) + 0.1mM 2-mercapto-ethanol (Thermo Fisher #31350010) + 1% L-Glutamine (Eurclone #ECB3000D) + 1% penicillin/streptomycin (Euroclone #ECB3001D) + 0.02mM Folic Acid (Sigma #I7508) + 0.2mM Myo-inositol (Sigma #I7508); and freshly added with 200U/ml recombinant human IL-2 (Prepotech , #200-02) according to Manufacturer’s instructions [69]. MDA-MB-436 were maintained in adhesion cell cultures and grown in 44.5% Dulbecco’s Modified Eagle Medium (DMEM), GlutaMAXTM (Thermo Fisher, #61965026) + 44.5% Ham’s F-12 Nutrient Mix, GlutaMAX™ (Thermo Fisher, #31765035) + 10% FBS South America + 1% penicillin/streptomycin. All cell lines were grown in an incubator at 37°C with a humidified atmosphere at 5% CO_2_, regularly tested for mycoplasma, and passed once 70-90% of confluence was reached. For sample preparation, NK92 cells were harvested, centrifuged for 5min at 1500rpm, washed with 10mL PBS, and resuspended in their culture medium for the subsequential cell counting. MDA-MB-436 cells were detached following standard procedure, meaning medium removal, PBS washing, and incubation with Trypsin-EDTA (Sigma #T8154) for 5 min at 37°C. Next, the cell culture medium was added to block trypsin activity. Cells were counted using Burker Cell Counter and Trypan Blue dye (Sigma #T8154) for viability assessment. Two independent counts were performed for each cell line by diluting 1:2 cell suspension with Trypan Blue, and 0.01mL was loaded in the cell counter. The experimental cell suspension was prepared considering the average live cell count, and each cell line was diluted in its culture medium to 2×10^5^ cells/mL of which 200μL were injected into the microfluidic channel.

### 2.2 Holographic dataset creation

To perform holographic recording of flowing cells, we used a HIFC system equipped with a digital holographic microscope employing a Mach Zehnder interferometer based on off-axis configuration coupled with a microfluidic channel (MC, Microfluidic ChipShop 10 000 107 – 200 μm × 1000 μm × 58.5 mm), as sketched in Fig. 1(a) and described in [21]. The interference fringe pattern is generated using a beam splitter cube (BS) that recombines the object and reference beams. These two beams originate from the division of a laser-emitted light wave (Laser Quantum Torus 532, beam diameter 1.7 ± 0.2 mm) by means of a polarizing beam splitter (PBS). Two half-wave plates (HWPs), positioned before and after the PBS, are used to adjust the relative intensities of the beams while preserving their polarization states. The object beam illuminates the cells as they undergo combined rotational and translational motion within the MC. The light scattered by the cells is collected by a microscope objective (MO1, Zeiss Plan-Apochromat 40×, NA = 1.3, oil immersion) and subsequently relayed through a tube lens (L1). In parallel, the reference beam is expanded by a beam expander (BE) composed of a second microscope objective (MO2), a second tube lens (L2), and a pinhole. The resulting interference pattern (hologram) is recorded by a CMOS camera (Genie Nano-CXP, 5120×5120 pixels with 4.5 μm pixel size) operating at a frame rate of 30 Hz and an exposure time of 50 μs. Cells are introduced into the MC using an automated syringe pump (CETONI neMESYS 290N), which delivers the sample at a controlled flow rate of 75 nL/s.

**Figure 1.**
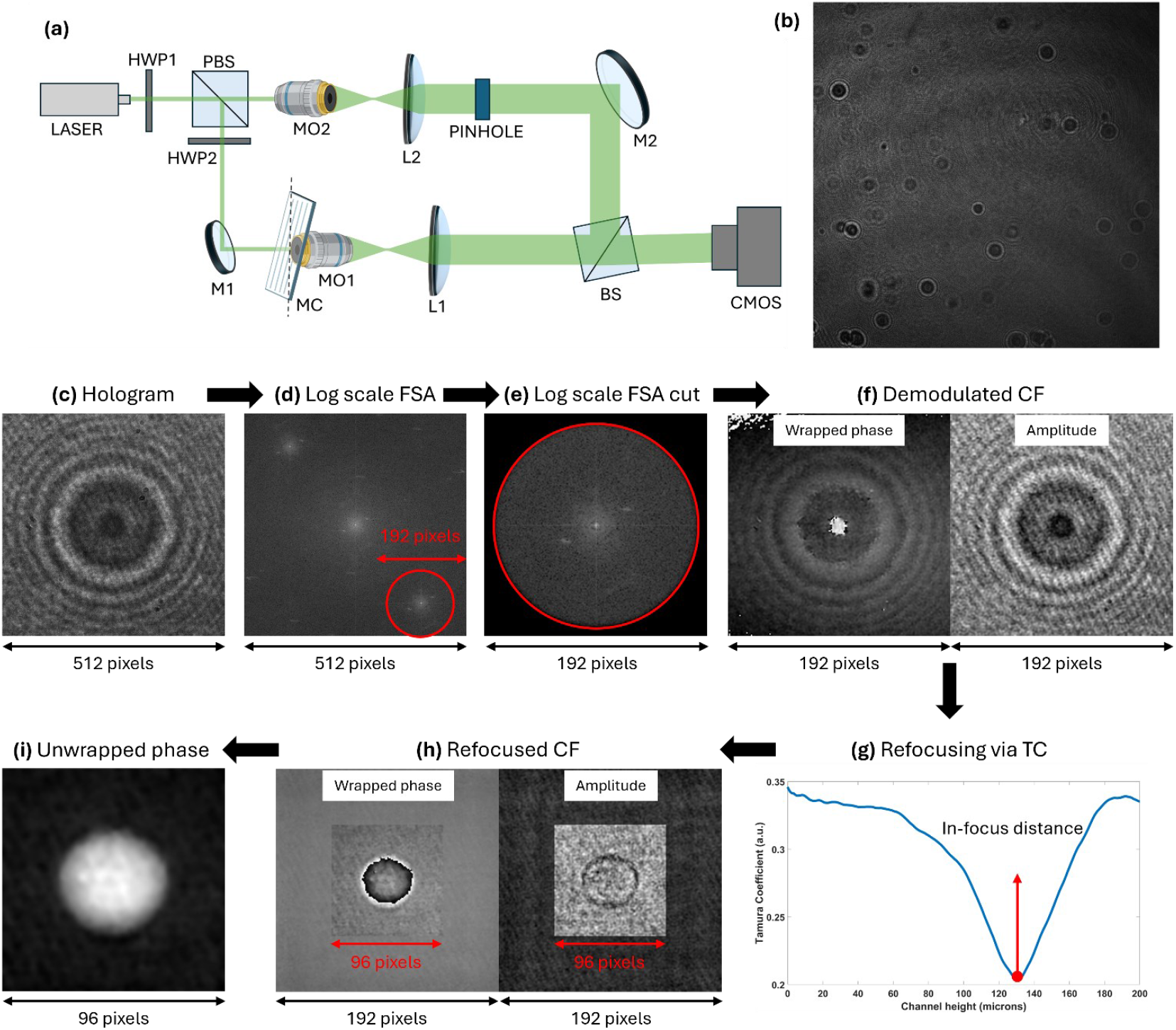
HIFC experiments and numerical processing. **(a)** Sketch of the HIFC experimental system. PBS – polarizing beam splitter; HWP – half wave plate; M – mirror; MO – microscope objective; P – pinhole; L – lens; BS – beam splitter; MC – microfluidic channel; CMOS – camera. **(b)** Example of recorded digital hologram. **(c-i)** Stages of the holographic reconstruction processing from the holographic ROI to the unwrapped phase image.

An example of 5120×5120 digital hologram recorded by the HIFC system is shown in Fig. 1(b). Cells are detected and tracked from the holographic video sequence recorded by the HIFC system. In each frame, a region of interest (ROI) of sizes 512×512 is selected around each detected cell, as displayed in Fig. 1(c). Therefore, the holographic video sequence is converted into multiple sequences of holographic ROIs, each of which contains a single snapshot of a cell. A phase image is then computed from each holographic ROI using the optimized reconstruction pipeline inspired by the methodology reported in [70]. By applying a circular band-pass filtering in the Fourier domain (Fig. 1(d)), the real diffraction order is numerically filtered out and a cut around the circular filter is applied (Fig. 1(e)), thus obtaining the demodulated CF (complex-valued image) of sizes 192×192 (Fig. 1(f)) without losing information, as demonstrated in [70]. Then, through the minimization of the Tamura coefficient (Fig. 1(g)), the proper focus distance is evaluated in order to achieve the numerical refocusing by propagating the demodulated CF at such distance using the angular spectrum method. Basically, the refocused CF minimizes the diffraction rings around the cells, hence we can further reduce the size of the complex-valued image up to 96×96 without losing information (Fig. 1(h)). Finally, the angular part of the refocused CF corresponds to the wrapped phase image on which the unwrapping algorithm based on least square minimization [71] is used to recover the real-valued 96×96 unwrapped phase image (Fig. 1(i)). Notice that, if needed, some additional reconstruction stages can be considered when residual phase aberrations occur or highly noisy data are recorded. In our reconstruction pipeline, polynomial fitting is used to remove the potential residual phase aberration just after the refocusing stage, while the WF2F denoising algorithm [72] is used to clean up phase jumps visibility just before the phase unwrapping stage.

Using the HIFC experimental system, we collected two datasets of raw holograms of sizes 5120×5120 containing NK92 and MDA-MB-436 cell lines, from which we extracted a total of 1877 single-cell holographic ROIs of sizes 512×512. Then, for each holographic ROI, we performed the reconstruction pipeline reported in Fig. 1(c-i), thus obtaining the datasets of demodulated CFs, refocused CFs, and unwrapped phase images. Each dataset was divided into training and test sets, as summarized in Table 1. Note that all holograms and their corresponding image representations in the test sets were collected in experiments separate from those used for the training sets.

**Table 1.** Number of single-cell images in the dataset for CNN classification across reconstruction stages.

| Cell Line | Training Set | Test Set |
| --- | --- | --- |
| MDA-MB-436 | 931 | 84 |
| NK92 | 846 | 76 |

## 3. Results and Discussion

The classification task for distinguishing MDA-MB-436 and NK92 cells is implemented using a generalized version of the TMEnet proposed in [21], which has demonstrated superior efficiency and accuracy compared to commonly used deep learning models. TMEnet follows a VGG-style architecture with eight convolutional layers (3×3), each followed by batch normalization and ReLU activation, and a max pooling layer (2×2, stride 2) after every two convolutions. The convolutional blocks are followed by a convolutional layer with a unitary-sized kernel and a global max pooling layer. A final ReLU activation and a dropout layer (p = 0.5) complete the architecture. The generalized model is obtained by parametrizing the number of filters in the convolutional layers using the variable K: the first four convolutional layers use K filters, the next four use 2K filters, and the final convolutional layer with unitary-sized kernel uses 16K filters. A fully connected layer maps to the number of classes (two in this case), and a softmax layer is used for classification, with weights optimized using cross-entropy loss. The TMEnet architecture is illustrated in Fig. 2, highlighting its ability to accept different input sizes: 512×512×1 raw holograms, 192×192×2 demodulated CFs, 96×96×2 refocused CFs, and 96×96×1 unwrapped phase images. For complex-valued inputs, the two channels represent the real and imaginary parts of the CF. For a fair comparison among TMEnets results, in all cases the training step was performed over 50 epochs with a batch size of 64, with a stopping criterion based on the best validation loss. We used the ADAM optimizer with a learning rate of 10^−4^. Training was performed in MATLAB^®^ R2025a on a workstation (Lenovo ThinkStation P5) equipped with an Intel^®^ Xeon^®^ W7-2475X CPU (2.59 GHz), 128 GB RAM, and an NVIDIA RTX 4000 Ada Generation GPU with 20 GB memory.

**Figure 2.**
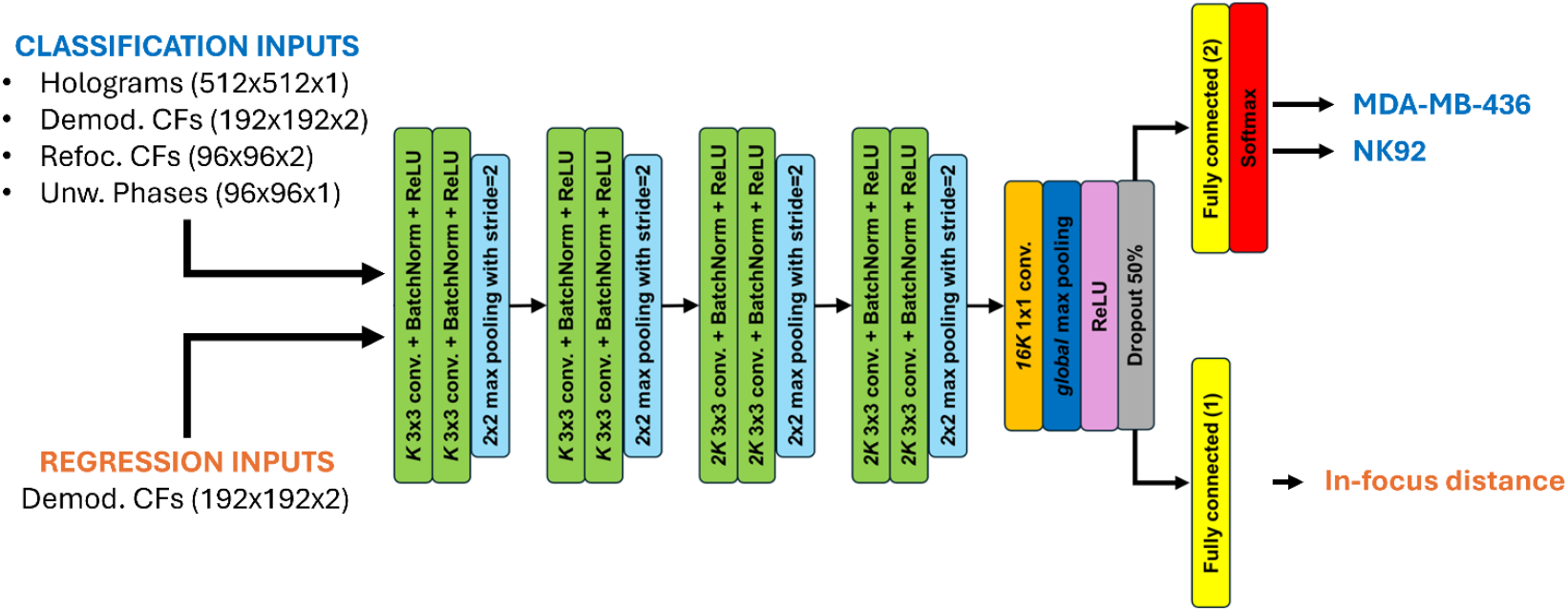
Generalized TMEnet. The parameter K corresponds to the number of filters in the first convolutional layer.

Table 2 summarizes the training and testing results of the generalized TMEnet with K=64, reporting image sizes, processing time per stage, cumulative reconstruction time, deployment time for class prediction, total classification time (reconstruction + deployment), maximum processable images per second (IPS), and classification accuracies on both validation and test sets. Each model is trained five times using different validation subsets, each comprising 10% of the training set, forming a 5-fold cross-validation scheme. The five trained networks are then evaluated on the same test set (Table 1), and the reported accuracies in Table 2 correspond to the average results across the five folds. In terms of performance, classification accuracy increases along the reconstruction stages, with the highest result achieved using unwrapped phase images, reaching an average accuracy of 95.87%, but at the cost of the highest computational load. Specifically, processing a raw hologram to recover the corresponding unwrapped phase image takes 0.4596 s, and the deployment time for class prediction is 0.0047 s, resulting in a maximum throughput of only ∼2 IPS. In contrast, the fastest computation is obtained when classifying demodulated CFs, achieving a potential throughput of 34 IPS with an average accuracy of 90.12%. Interestingly, this is faster than directly classifying raw holograms. The optimized demodulation stage [70] allows reducing the input size from 512×512×1 to 192×192×2, almost doubling the speed (from 18 IPS to 34 IPS) while slightly improving accuracy. The above results demonstrate the expected trade-off between accuracy and computing time, motivating the exploration of solutions that accelerate processing with only a minor reduction in accuracy.

**Table 2.** Performance of the trained TMEnets across reconstruction stages in terms of both computing time and average classification accuracies.

|  | Holograms | Demodulated CFs | Refocused CFs | Unwrapped Phases |
| --- | --- | --- | --- | --- |
| **Sizes** | $512 \times 512 \times 1$ | $192 \times 192 \times 2$ | $96 \times 96 \times 2$ | $96 \times 96 \times 1$ |
| **Single Stage Processing Time (sec)** | --- | 0.0177 | 0.3975 | 0.0444 |
| **Reconstruction Time (sec)** | --- | 0.0177 | 0.4152 | 0.4596 |
| **Deploy Time (sec)** | 0.0527 | 0.0112 | 0.0053 | 0.0047 |
| **Classification Time (sec)** | 0.0527 | 0.0289 | 0.4205 | 0.4643 |
| **Images Per Second (IPS)** | 18 | 34 | 2 | 2 |
| **Classification Accuracy Validation (over 5 trials)** | 86.29% | 87.30% | 91.91% | 93.15% |
| **Classification Accuracy Test (over 5 trials)** | 89.62% | 90.12% | 93.87% | 95.87% |

Analysis of the processing times indicates that the refocusing stage is the main bottleneck, as it is the slowest step due to the conventional iterative procedure required to recover the in-focus distance. To address this, deep learning-based algorithms can be employed to predict the in-focus distance, as demonstrated in [65–68]. Here, we propose to reuse TMEnet, suitably adapted for a continuous-valued regression problem. The network architecture is obtained from the corresponding classification layout by removing the last two layers and replacing them with a fully connected layer with a single-valued output. The model weights are optimized using a loss function based on the mean absolute error (MAE). Our implementation is shown in the bottom part of Fig. 2 using K=32, where the input is the demodulated CF and the output is the predicted in-focus distance. The dataset for training is created by merging the training sets of demodulated CFs using for the classification task, i.e. by considering all images from both NK92 and MDA-MB-436 cells in Table 1. The corresponding in-focus distances (i.e. ground truth data) are obtained by numerical refocusing using the Tamura coefficient, as described in Section 2.2. In this scenario, the reconstruction pipeline consists of demodulation, prediction of the in-focus distance, reconstruction of the refocused CF, and calculation of the unwrapped phase image, which is then fed to the classification network. With this approach, computation of the refocused CF is over 45 times faster than the conventional iterative process based on TC minimization.

Alternatively, a second solution involves training a CNN for end-to-end reconstruction, as demonstrated in [58–64]. Here, we implement a modified UNET architecture to predict a 96×96 unwrapped phase image directly from a 512×512 recorded hologram. Also in this case, the dataset for training is created by merging the training sets used for the classification task, i.e. all raw holograms and all unwrapped phase images from both NK92 and MDA-MB-436 training data in Table 1. The proposed architecture, illustrated in Fig. 3, is inspired by the well-established UNET with skip connections. Specifically, we introduce a correction in the decoder stage by adjusting the size of the first upsampling layer immediately after the latent space. The skip connections are also adapted to accommodate the change in feature map sizes. Also in this case we use MAE as the loss function. With this approach, the reconstruction process is accelerated by more than tenfold.

**Figure 3.**
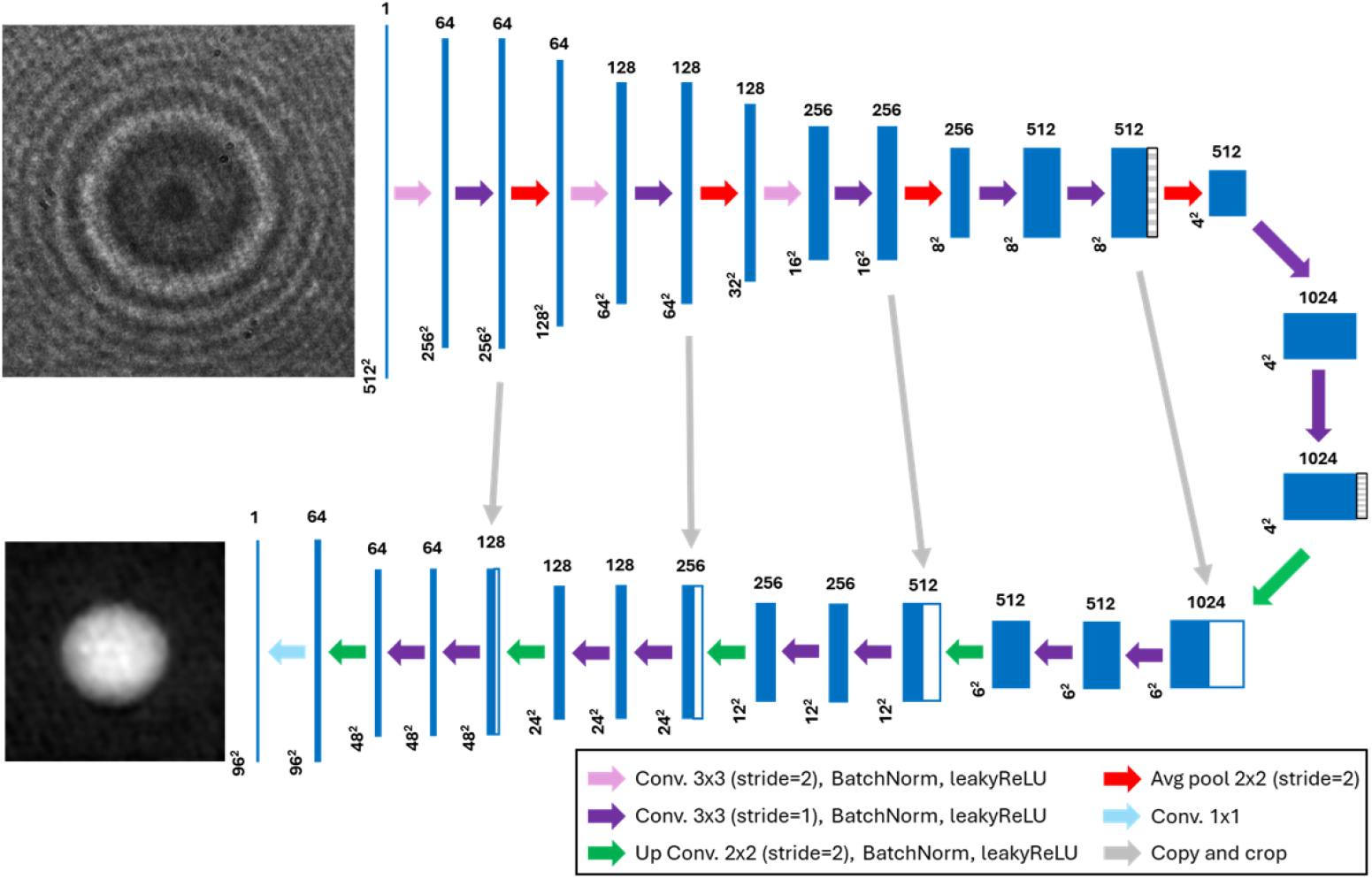
Modified UNET for end-to-end reconstruction. The network reconstructs the phase image directly from the holographic ROI by avoiding any holographic reconstruction stage.

The results of this analysis are reported in Table 3. To evaluate the performance of the alternative solutions, we use them to predict phase images of the test set, performing the full reconstruction process and measuring the corresponding classification times and IPS. The TMEnets trained on unwrapped phase images are then employed to compute the average classification accuracy across the five test trials, as in Table 2. Again, the results show a clear trade-off between speed and accuracy: the end-to-end UNET is 1.6× faster than the reconstruction pipeline based on the refocusing TMEnet, but with a 3% reduction in classification accuracy. This decrease is likely due to the lower quality of the phase images predicted by the UNET model. To investigate this aspect, we compute two fidelity metrics between predicted and reconstructed phase images, namely the Normalized Root Mean Square Error (NRMSE) and the Structural Similarity Index Measure (SSIM). In both cases, the phase images obtained using the refocusing TMEnet are closer to the unwrapped phase images reconstructed through the standard holographic processing pipeline, confirming the observed trade-off.

**Table 3.** Performance of the hybrid solutions based on CNN models to replace the automatic refocusing step or to perform end-to-end phase reconstruction.

|  | Refocusing TMEnet | End-to-End UNET |
| --- | --- | --- |
| **NRMSE** | 0.0205 | 0.0289 |
| **SSIM** | 0.9363 | 0.9119 |
| **Classification Time (sec)** | 0.0757 | 0.0461 |
| **Images Per Second (IPS)** | 13 | 21 |
| **Classification Accuracy Test (over 5 trials)** | 94.25% | 91.25% |

In summary, we identify six possible strategies to perform the classification task from which the optimal one needs to be selected. To do this, we perform a Pareto analysis of them, as shown in Fig. 4. A strategy is considered Pareto-optimal if no other strategy is simultaneously more accurate and faster [73]. The resulting Pareto front highlights the set of non-dominated strategies, providing a clear visualization of the best compromises between performance and efficiency. Non-dominated strategies are marked in green in Fig. 4, with the Pareto front connected by a black line. These include the use of demodulated CFs and unwrapped phase images, as well as hybrid solutions such as the refocusing TMEnet and the end-to-end UNET. Strategies outside the Pareto front are dominated, representing non-optimal trade-offs since they are outperformed in both metrics by at least one alternative. Dominated strategies are marked in red in Fig. 4 and include the use of raw holograms, which are outperformed by demodulated CFs, and refocused CFs, which are outperformed by the refocusing TMEnet. Among the remaining four strategies, the best trade-off is identified by the highest slope of the Pareto front, corresponding to the Refocusing TMEnet approach. Therefore, the optimal solution for the proposed classification task, simultaneously maximizing classification accuracy and minimizing classification time, consists of the following pipeline: raw hologram demodulation, prediction of the in-focus distance using the Refocusing TMEnet, reconstruction of the refocused CF using the predicted distance, and computation of the unwrapped phase image. The resulting phase image is then fed into the trained TMEnet classification model to predict the cell population.

**Figure 4.**
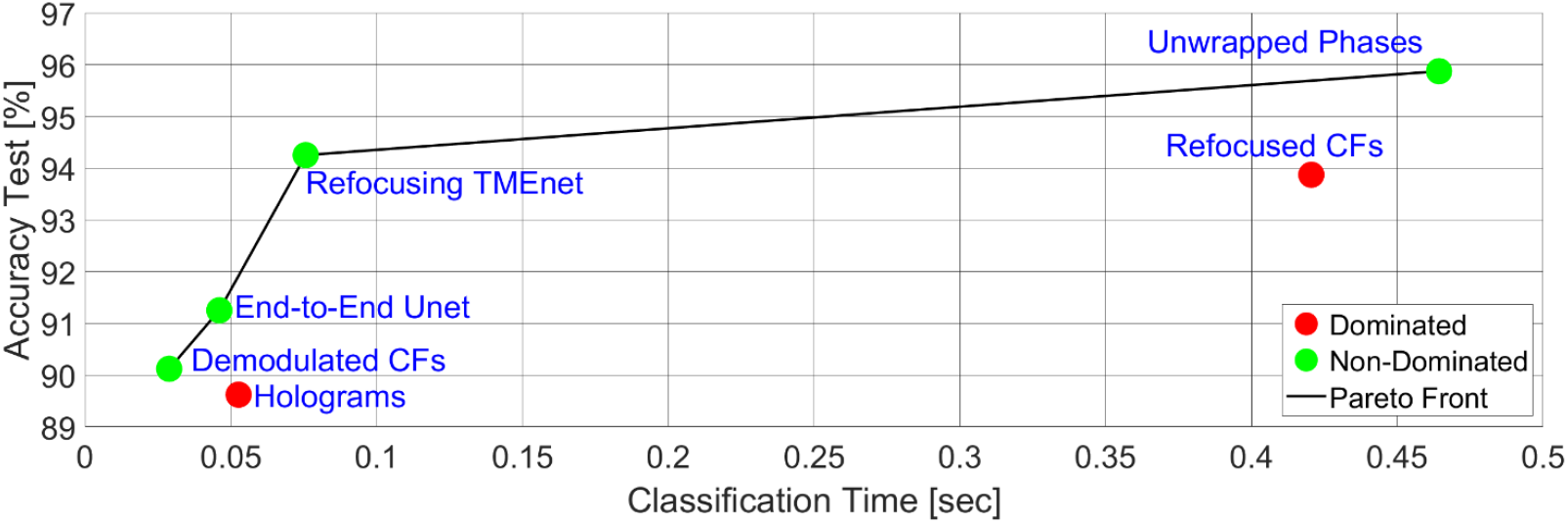
Comparison between different classification strategies in terms of accuracy over the test set and classification time. Dominated strategies are marked in red, while non-dominated strategies within the Pareto front (black line) are marked in green.

## 4. Conclusion

In this paper, we have systematically investigated the impact of image representation on deep learning-based single-cell classification in HIFC, explicitly quantifying the trade-off between classification accuracy and computational efficiency across successive holographic reconstruction stages. First, our study provided a structured comparison between the main four image representations within the holographic reconstruction pipeline, i.e. the direct classification of raw holograms, demodulated CFs, refocused CFs, and unwrapped phase images. For this analysis, a generalized version of the TMEnet model proposed in [21] was adapted to be trained with input having different sizes in order to perform a fair comparison among the four classification schemes. By benchmarking these strategies, we demonstrated that classification accuracy increases monotonically with reconstruction completeness. The highest performance was achieved using fully reconstructed unwrapped phase images, confirming that quantitative phase information encodes discriminative biophysical features essential for reliable cell-type identification. However, this gain in accuracy comes at the expense of substantial computational cost, primarily dominated by the iterative refocusing stage. Conversely, simplified representations such as demodulated complex fields offer a dramatic increase in throughput with only moderate accuracy degradation. Interestingly, demodulation not only reduces input dimensionality but also improves predictive performance compared to raw holograms, indicating that partial physics-based preprocessing can enhance learning efficiency by suppressing potential experimental artifacts and biases while preserving informative content.

To address the computational bottleneck introduced by the holographic processing pipeline, we explored two hybrid strategies exploiting deep learning–based acceleration paradigms: (i) a regression-based TMEnet for in-focus distance prediction to replace the slowest reconstruction step and (ii) a modified UNET for end-to-end phase reconstruction. Both approaches significantly reduced processing time, with the regression-based refocusing strategy emerging as the most favorable compromise between classification speed and accuracy. Indeed, the Pareto analysis provided a rigorous multi-objective evaluation, identifying non-dominated solutions and the optimal solution able to simultaneously maximize the accuracy and minimize the computational burden. In particular, replacing only the refocusing stage with a learned TMEnet predictor preserved both high phase fidelity and classification accuracy while increasing throughput by more than one order of magnitude compared to the conventional pipeline.

From a broader perspective, these findings underline that image representation is not a neutral preprocessing choice but a central design variable. Of course, the optimal strategy depends on the specific application constraints: high-throughput screening may prioritize demodulated representations, whereas diagnostic scenarios demanding maximal accuracy may justify full phase reconstruction. Hybrid strategies offer a flexible intermediate solution, enabling adaptive deployment based on available hardware resources and latency requirements.

The present study focuses on a binary classification task, involving NK92 and MDA-MB-436 cells. The importance of NK fast enumeration may be useful while assessing the response to both chemotherapy and immunotherapy-based protocols. Indeed, the correct quantification of these immune cells in the tumor microenvironment currently faces several technical challenges due to the general low infiltration rate and the absence of a standardized methodology among Pathologists. While only this proof-of-concept case study is considered, the proposed framework is general and can be extended to multi-class problems, unbalanced datasets, or other biophysical phenotyping tasks. Future work may investigate domain adaptation strategies, uncertainty quantification, hardware-aware model compression, and real-time edge deployment to further enhance translational potential. Additionally, integrating temporal dynamics from holographic video sequences could provide complementary information beyond single-frame representations.

## Data availability statement

The data that support the findings of this study are available upon reasonable request from the authors.

## Acknowledgements

This work was supported by project PRIN 2022, Computationally aided Opto-mechano-fluidic pLatform for Label-free intelligent tumor microEnvironment Cell sorTing (COLLECT) Prot. 202275PJRP, funded by the Italian Ministry of University & Research in the framework of the European Union program Next Generation EU, CUP: B53D23002280006.

## Author contributions

Daniele Pirone: Formal analysis, Software, Visualization, Writing – original draft. Beatrice Cavina: Resources. Giusy Giugliano: Investigation. Francesca Nanetti: Resources. Francesca Reggiani: Writing - Review & Editing, Resources. Lisa Miccio: Investigation, Validation. Ivana Kurelac: Supervision, Writing - Review & Editing, Resources. Pietro Ferraro: Conceptualization, Writing – original draft. Pasquale Memmolo: Conceptualization, Methodology, Supervision, Writing – original draft, Software, Funding acquisition.

